# Synteny-aware functional annotation of bacteriophage genomes with Phynteny

**DOI:** 10.1101/2025.07.28.667340

**Authors:** Susanna R. Grigson, George Bouras, Bhavya Papudeshi, Vijini Mallawaarachchi, Michael R. Roach, Przemyslaw Decewicz, Robert A. Edwards

**Affiliations:** Flinders Accelerator for Microbiome Exploration, College of Science and Engineering, Flinders University, Adelaide, SA, 5042, Australia; Adelaide Medical School, Faculty of Health and Medical Sciences, The University of Adelaide, Adelaide, SA, 5005, Australia; The Department of Surgery - Otolaryngology Head and Neck Surgery, University of Adelaide and the Basil Hetzel Institute for Translational Health Research, Central Adelaide Local Health Network, Adelaide, SA, 5005, Australia; Flinders Health and Medical Research Institute, College of Medicine and Public Health, Flinders University, Adelaide, SA, 5042, Australia; Department of Environmental Microbiology and Biotechnology, Institute of Microbiology, Faculty of Biology, University of Warsaw, Warsaw, Poland

## Abstract

Accurate genome annotation is fundamental to decoding viral diversity and understanding bacteriophage biology; yet, the majority of bacteriophage genes remain functionally uncharacterised. Bacteriophage genomes often exhibit conserved gene order, or synteny, that reflects underlying constraints in genome architecture and expression. Here, we present *Phynteny*, a genome-scale, deep learning framework that leverages gene synteny to predict the function of unknown bacteriophage genes. Phynteny integrates protein language model embeddings with positional encoding, bidirectional long short-term memory, and transformer encoders featuring circular attention to learn genome-wide organisational patterns. Trained on a dereplicated dataset of over 280,000 bacteriophage genomes, Phynteny achieves high predictive performance (AUC > 0.84) across the nine PHROG functional categories and confidently assigns putative functions to improve the number of annotated genes in phage isolate genomes by 14%. To assess the validity of these predictions, we compared them with annotations derived independently using protein structural information, revealing broad functional concordance and additional confidence in Phynteny predictions. By incorporating genomic context into functional annotation, Phynteny offers a novel approach to illuminate the functional landscape of viral dark matter and is available at https://github.com/susiegriggo/Phynteny_transformer.

## Introduction

Bacteriophages (phages), viruses that infect bacteria and replicate using their host’s metabolism, are the most abundant and diverse biological entities on Earth^1–3^. Found in every known ecosystem, phages shape microbial evolution through predation and horizontal gene transfer. As key regulators of microbial communities, phages mediate genetic exchange, drive bacterial population dynamics, and influence biogeochemical cycling^4–6^. Growing interest in phage therapy, using phages to target and eliminate pathogenic bacteria, has further highlighted their potential as alternatives to antibiotics in the fight against multidrug-resistant infections^7,8^.

Despite the rapid expansion of viral sequence databases^9–12^, functional annotation of phage genomes remains a significant challenge. Viral gene functions are routinely inferred using sequence homology, where functional labels are transferred to unknown genes from homologous proteins of known function^13^. This approach often uses alignment-based methods, such as profile Hidden Markov Models, as implemented in tools like Pharokka^14^ and geNomad^15^. However, the vast sequence diversity of phages and limited representation in reference databases severely constrain homology-based predictions. As a result, 60–80% of phage genes lack any functional annotation^16–18^. This persistent sequence–function gap hinders our understanding of viral biology and poses potential risks for phage therapy, where the inadvertent transfer of genes encoding toxins or virulence factors remains a significant concern^19,20^. Improving functional annotation strategies requires new approaches beyond sequence similarity to enable broader, more accurate functional annotation.

Gene annotation frameworks have been devised to characterise the functional repertoire of bacteriophage genes^21^. Among these, Prokaryotic Virus Remote Homologous Groups (PHROGs) define homologous gene families and assign them to one of nine functional categories^22^. These categories encompass structural proteins that form the phage virion, replication and transcription machinery, elements involved in host integration and lysis, and additional auxiliary functions. However, most PHROGs have undetermined functions and only 13% (5,133 of 38,880) are linked to a defined category. To address this challenge, machine learning approaches have emerged that capture functional information beyond what is detected with alignment-based methods. Phage Artificial Neural Networks (PhANNs)^23^ uses neural networks trained on numerical representations of amino acid composition, sequence motifs, and physicochemical properties to infer gene function. Viral Protein Function prediction using Protein Language Model (VPF-PLM)^24^, utilises numerical vectors generated from protein language models pre-trained on billions of protein sequences to classify phage gene functions. Alternatively, Phold (https://github.com/gbouras13/phold) uses protein structural information to generate functional annotations using the ProstT5 protein language model^25^. While these approaches provide an alternative to sequence homology for inferring phage gene functions, these models treat each gene in isolation, overlooking the broader genomic contexts that often inform function.

Phage genomes exhibit highly conserved gene order, or synteny, arising from evolutionary constraints associated with viral replication, assembly, and infection cycles^26,27^. This results in distinct, ordered modules of genes with specific functions ^28,29^. Structural genes encoding tail proteins, capsid proteins, baseplates, and other virion components are among the most highly conserved in their order, reflecting constraints linked to assembly and function. Other modules govern integration into the host genome, DNA replication and repair, transcriptional regulation, and host cell lysis. This modular architecture suggests that gene function may be inferred not only from sequence similarity but also from a gene’s relative genomic position^30^. In bacterial genomics, gene order has been successfully leveraged to predict gene function across operons^31^ and metagenomic contigs^32,33^. However, for phage genome annotation, this approach remains underutilised, as current annotation methods do not capture the interdependencies and long-range relationships between genes. We propose that integrating syntenic information with protein language model embeddings and homology-based annotations can enhance phage functional annotation.

Here, we present *Phynteny*, a transformer-based model for phage gene functional annotation that integrates genomic context and protein-level information to infer function across viral genomes. By encoding phage genomes as ordered matrices of gene-level features, including protein language model embeddings, gene orientation, length and functional annotations, our model learns conserved patterns of gene synteny that underpin viral genome architecture. Trained on a dereplicated and quality-filtered subset of over 289,000 genomes from the PhageScope^9^ database, Phynteny accurately predicts gene function across nine PHROG functional categories, including genes with no detectable homology to known proteins. This approach marks a shift toward context-aware genome annotation, bridging local protein features with genome-scale patterns. Our results show that incorporating gene order significantly improves the rate of phage genome annotation and reveals underlying organisational principles of phage genomes.

## Results

### Constructing a Transformer model for synteny-aware functional annotation

To leverage the conserved gene-ordering properties of phage genomes for functional annotation, we developed *Phynteny*—a genome-scale classification model that predicts the function of unannotated phage genes using syntenic context. The model assigns genes to one of nine PHROG functional categories, representing core viral processes, based on their position within the genome and surrounding gene features. To train the model across the broad diversity of bacteriophage genomes, we curated a non-redundant, high-quality dataset comprising 289,106 genomes from the PhageScope database^9^, applying stringent filtering criteria to exclude incomplete or low-confidence assemblies (Fig. 1A). Across these genomes, an average of 29% of genes were annotated with PHROGs, with the proportion varying by category: head and packaging (6.5%), connector (1.5%), tail (5.0%), lysis (1.8%), transcription (1.8%), DNA, RNA and nucleotide metabolism (nucleic acid metabolism) (8.0%), moron, auxiliary metabolic gene and host takeover (morons) (1.0%), transcription regulation (1.8%), and hypothetical proteins (71%)(Fig. S1)

**Figure 1.**
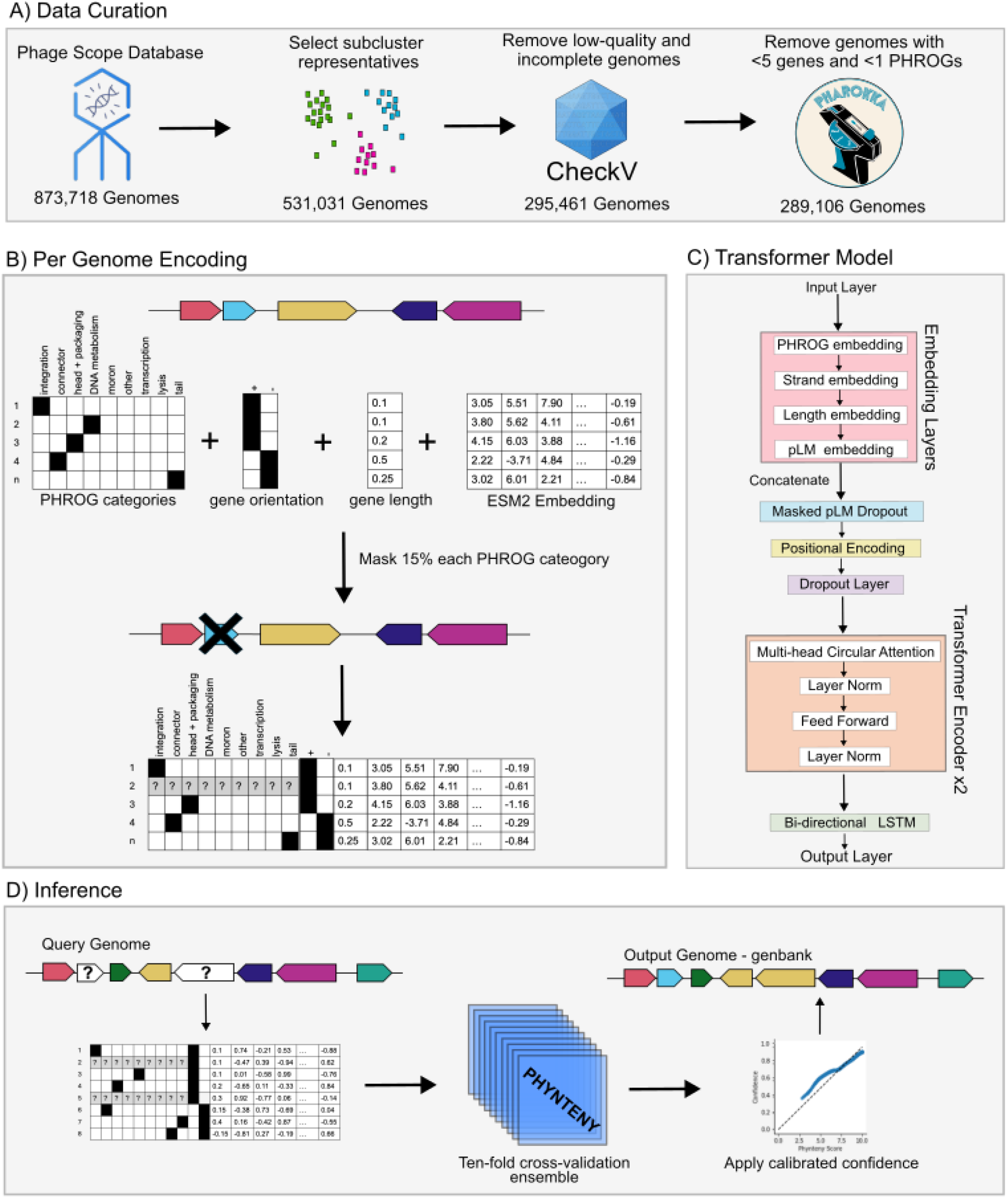
Overview of training synteny-aware protein function classifier and the Phynteny-transformer annotation tool. **A**. Curation of data used for model training. One genome was selected per PhageScope subcluster. Genomes flagged as low or undetermined quality by CheckV, or containing fewer than five genes or lacking PHROG annotations assigned by Pharokka (based on the PHROG database), were excluded. **B**. Feature encoding for input representation. For each genome, gene feature vectors were constructed by concatenating one-hot encoded functional annotations, strand orientation, gene length, and protein language model embeddings (ESM-2). To enable supervised learning, a subset of 15% of genes from each PHROG category was randomly masked as ‘unknown’. **C**. Architecture of the Phynteny transformer model. Each input feature type is first embedded via a dedicated trainable layer. These are followed by dropout, positional encoding, and BiLSTM layers. A circular multi-head self-attention mechanism is applied within the transformer encoder to account for genome circularity and synteny. **D**. Inference workflow for phage gene function prediction. Encoded genome features are passed through an ensemble for ten independently trained models obtained via cross-validation. Model outputs are calibrated and provide predictions and their confidence to the user in an updated GenBank form.

Phynteny encodes each genome as a matrix, with rows corresponding to genes in their native genomic order. For each gene, a feature vector is constructed comprising one-hot encoded PHROG annotations, strand orientation, gene length, and protein embeddings obtained from the ESM2 protein language model (ESM2-50M)^34,35^. In each training pass, a dynamic masking procedure is applied where 15% of the genes from each functional category are randomly designated as ‘unknown’. This masking probability is adjusted using inverse frequency weighting, where rare functional categories receive higher masking probabilities to compensate for their underrepresentation in the training data, while common categories receive correspondingly lower probability (See Methods). This proportion is commonly used in masked language modeling frameworks, including BERT and related transformer-based models, as it provides a balance between learning efficiency and context retention^34,36^. These masked genes serve as prediction targets, enabling supervised learning by computing the loss relative to their known PHROG categories (Fig. 1B).

Phage genome matrices are processed by a deep-learning architecture that captures higher–order dependencies in gene arrangement. Input features are first transformed through individual embedding layers, which convert the inputs to dense vector representations. To discourage over-reliance on the pre-trained protein embeddings and promote the use of genomic context, a dropout layer^37^ is applied selectively to the protein language model embeddings of masked genes. Spatial relationships among genes are preserved through sinusoidal positional encodings, while a bidirectional long short-term memory layer (BiLSTM) captures context from both upstream and downstream elements. Two circular self-attention transformer layers^38^ follow, employing relative position attention mechanisms that account for the natural circularity of phage genomes. A final fully connected output layer projects the learned representations to the nine PHROG categories (Fig. 1C).

We developed a command-line tool that applies the trained ensemble to predict the function of unknown genes in annotated phage genomes. During inference, Phynteny takes as input GenBank files annotated with PHROG categories, such as those produced by Pharokka^14^, where each coding sequence includes a PHROG label as the ‘function’ qualifier. Predictions are generated using an ensemble of ten independently trained models from cross-validation. Each model produces an independent prediction, and the most confident output is selected for each unknown gene. Annotated genomes are returned as GenBank files and tab-separated (TSV) summary files, with each prediction accompanied by a confidence score, enabling seamless downstream analysis and integration with existing phage annotation workflows (Figure 1D). For advanced users, Phynteny also includes training scripts to build custom models from PHROG-annotated datasets.

### Phynteny utilises phage genomic synteny

To investigate patterns of gene organisation, we analysed a subset of phages from the curated PhageScope dataset that contained at least one gene from each of the following categories: ‘integration and excision’, ‘head and packaging’, ‘nucleic acid metabolism’, and ‘tail’, and had a genome length of <60 kb (26,091 genomes). Although not all phage genomes are truly circular, many being circularly permuted or terminally redundant rather than physically circular in the capsid, we treated them as circular to enable comparisons of gene order that are not biased by arbitrary genome start sites^13^. Clear trends in gene ordering emerged (Fig. 2A). Nucleic acid metabolism genes typically precede head and packaging genes, which are followed by tail and connector genes. In contrast, genes from the other and moron categories are more dispersed across genomes. Groups of phage genomes with similar modular gene arrangements emerged, indicative of conserved syntenic arrangements that transcend phage lineages.

**Figure 2.**
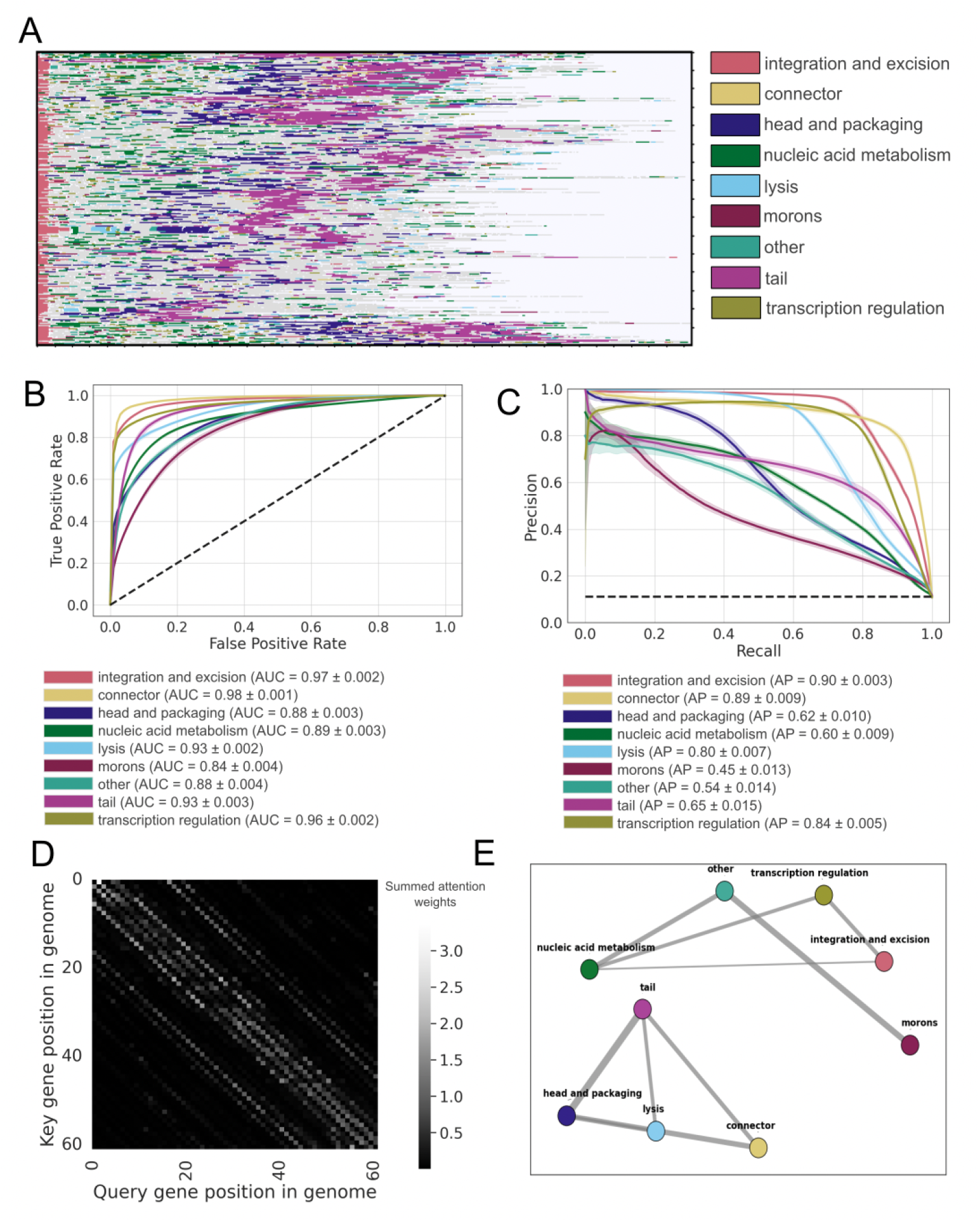
Transformer-based model captures gene-order-dependent signals for PHROG functional classification. **A**. Syntenic organisation of genomes from the curated PhageScope dataset. Genomes that contained at least one gene from the ‘head and packaging’, ‘integrase and excision’, ‘nucleic acid metabolism’ and ‘tail’ PHROG categories were selected (26,091 genomes). Genomes were recircularised such that the first gene corresponds to the ‘integration and excision’ category, and reoriented to ensure the first ‘head and packaging’ gene precedes the first ‘tail’ gene. Rows represent individual genomes and are clustered using the unweighted pair group method with arithmetic mean (UPGMA). **B**. Receiver operating characteristic (ROC) curves showing classification performance across PHROG categories, with area under the curve (AUC) and standard deviation calculated over 10-fold cross-validation. **C**. Precision Recall (PR) curves showing classification performance across PHROG categories, with average precision (AP) and standard deviation calculated over 10-fold cross-validation. **D**. Self-attention map from a representative genome (*Geobacillus* phage phiOH2; AB823818), showing the summed attention weights across all transformer layers of the first model in the Phynteny ensemble. The attention weights range from 0 (black) to 3.38 (white). The white lines near the diagonal show that the model is attending to neighbouring genes, while the more dispersed lighter points indicate the model is attending to distant genomic regions. The model attends both to neighbouring genes and distant genomic regions, reflecting context-aware information aggregation. **E**. Network representation of inter-category attention correlations. The network was constructed from data generated during the first fold of cross-validation using the corresponding model. Nodes represent PHROG functional categories, and edges reflect pairwise Pearson correlation coefficients (r > 0.5) between attention scores; edge width is proportional to correlation strength.

To determine whether the transformer-based Phynteny model learns PHROG functional categories, we performed 10-fold cross-validation to assess its performance on held-out data. The model effectively distinguished between functional categories, achieving an area under the ROC curve (AUC) greater than 0.84 and an average precision (AP) exceeding 0.45 across all PHROG categories (Fig. 2B–C). Structural genes (head and packaging, connector, and tail) and regulation genes (transcriptional regulation and lysis) achieved consistently high AUC and AP scores. In contrast, the moron category exhibited the lowest performance (AUC = 0.84, AP = 0.45), followed by the other category (AUC = 0.88, AP = 0.54), reflecting their more variable positioning and functional heterogeneity.

We analyzed the attention weights from Phynteny’s transformer layers to identify genomic features that contribute to its predictions beyond those encoded in protein language model embeddings. In an individual genome, attention signals appear both along the main diagonal and in off-diagonal regions of the gene-by-gene attention matrix. This indicates that Phynteny consistently attends to both adjacent genes and more distantly located regions of the genome, capturing local and long-range dependencies (Fig. 2D, S2). Relationships between functional categories were quantified by aggregating between-category attention scores and calculating Pearson correlation coefficients (Fig. 2E). Strong interdependencies were observed among structural modules, particularly tail, head and packaging, and lysis genes, indicating that the locations of these genes could be learned to predict the function of other genes. Similarly, genes associated with integration and excision, transcriptional regulation, moron auxiliary metabolic functions, and other accessory functions show a strong dependence on genes involved in DNA, RNA, and nucleotide metabolism (Fig. 2E).

### Gene-order enhanced functional annotation reaches viral dark matter

We applied Phynteny to a dataset of 16,442 previously unseen phage genomes identified as prophages within bacterial genomes^39^. Among the 1,026,848 proteins in this dataset, 195,364 (19.0%) had an assigned PHROG function. To estimate the confidence of Phynteny’s predictions for the remaining 831,484 proteins, raw output probabilities were calibrated using isotonic regression, enabling consistent interpretation of model certainty across functional categories (Fig. S3, Table S1). Connectors, integration and excision, head and packaging, and lysis genes exhibited a higher proportion of high-confidence predictions (score ≥ 0.8), whereas moron genes, despite being the most frequently predicted category, were predominantly assigned low confidence (Fig. 3A). Applying Phynteny’s default confidence threshold of 0.8 resulted in functional annotations for 342,226 proteins, an increase of 146,862 annotations compared to the initial set. This represents functional assignments for 14% of all prophage proteins, corresponding to a 75% increase over the number of proteins originally annotated (Fig. 3B).

**Figure 3.**
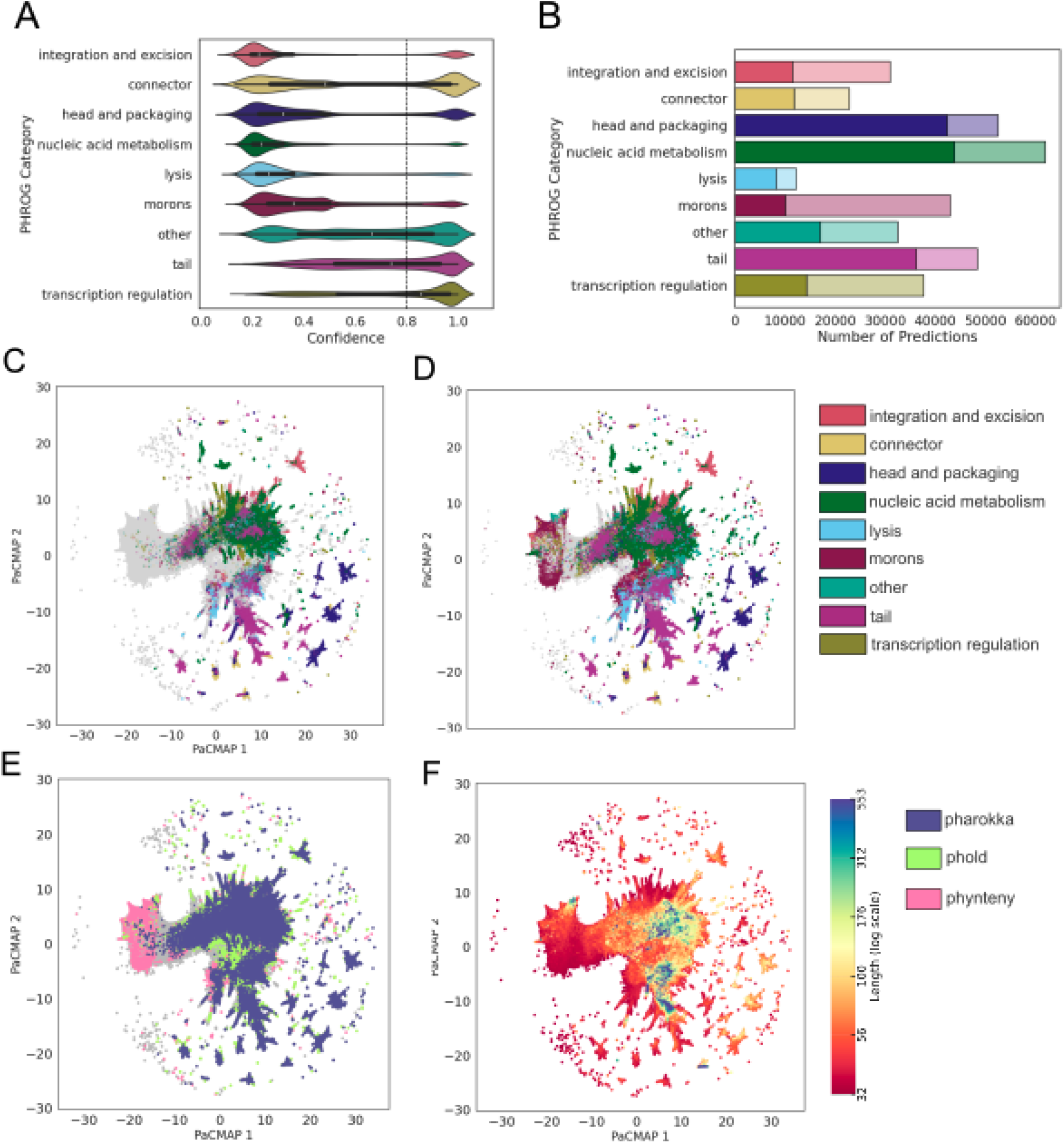
Phynteny illuminates phage dark matter. **A**. Confidence scores distribution for PHROG category predictions for sequences in the curated prophage dataset. Predictions were made for integration and excision (n=100,218), connector (n=28,557), head and packaging (n=52,139), nucleic acid metabolism (n=222,157), lysis (n=42,556), morons (n=274,445), other (n=41,179), tail (n=28,318) and transcription regulation (41,915). **B**. Number of annotations per PHROG category for genomes in the curated prophage dataset before (filled region) and after (shaded region) annotation with Phynteny. **C**. ESM2 embeddings of proteins in the curated prophage dataset visualised using pairwise controlled manifold approximation dimension reduction (PacMAP) (n_neighbours =20). Proteins are coloured by their PHROG category assigned using sequence homology via Pharokka. **D**. ESM2 embeddings of proteins in the curated prophage dataset visualised using PacMAP. Proteins are coloured by their PHROG category following annotation with Phynteny. **E**. ESM2 embeddings of proteins in the curated prophage dataset visualised using PacMAP. Proteins are coloured by which annotation method they could be annotated by (Pharokka, Phold, Phynteny). **F**. ESM2 embeddings of proteins in the curated prophage dataset visualised using PacMAP. Proteins are coloured by their length (amino acids).

For the 831,484 prophage proteins lacking prior functional annotations, Phynteny assigned functions to 113,755 proteins (13.6%) by linking them to PHROGs previously classified as “unknown,” spanning 9,481 such PHROGs. Following Phynteny annotation, 2,168 of these unknown PHROGs (36,671 proteins) were assigned to at least one known functional category. The largest number were assigned to morons (442 PHROGs), followed by nucleic acid metabolism (361), tail (306), other (276), transcription regulation (194), head and packaging (151), integration and excision (115), connector (115), and lysis (95) (Fig. S4). This reclassification demonstrates that Phynteny substantially reduces the proportion of uncharacterized PHROGs and expands the functional landscape of prophage genomes. Overlaps were observed between several categories, particularly among morons, nucleic acid metabolism, and tail proteins, suggesting functional diversity within individual PHROGs or potential multifunctionality of the encoded proteins.

Analysis of ESM2 embeddings revealed that proteins with prior homology-based annotations clustered within well-characterised regions of the embedding space (Fig. 3C). In contrast, many of the new Phynteny predictions extended into previously unannotated regions (Fig. 3D), capturing proteins overlooked by both homology-based tools (e.g. Pharokka) and structure-based approaches (e.g. Phold; Fig. 3E). Notably, a substantial fraction of these newly annotated proteins correspond to short sequences that commonly evade annoation by other methods (Fig 3F).

The Phynteny model combines both gene-order and protein embeddings. To isolate the contribution of gene-order modelling, we trained a simplified variant of Phynteny lacking the transformer and BiLSTM layers. This ablated model produced fewer high-confidence predictions across functional categories (Fig. S5). These results highlight the importance of modelling genomic context for improving generalisability and annotation coverage beyond categories dominated by homology signals.

### Gene-order enhanced annotation operates across phage families

To evaluate the lineage-specific benefits of gene order-informed annotation, we applied Phynteny to all isolate phage genomes in the INfrastructure for a PHAge Reference Database (INPHARED)^11^, which compiles all complete phage genomes and associated metadata from the National Center for Biotechnology Information’s GenBank database^40^. Genomes were first annotated using Pharokka, assigning functions based on sequence homology. Phynteny was then used to infer additional functions for unannotated genes based on genomic context. In individual genomes, Phynteny frequently assigned functions in a modular fashion, often filling in long contiguous regions with consistent functional categories (Fig. 4A), including across poorly annotated genomes.

**Figure 4.**
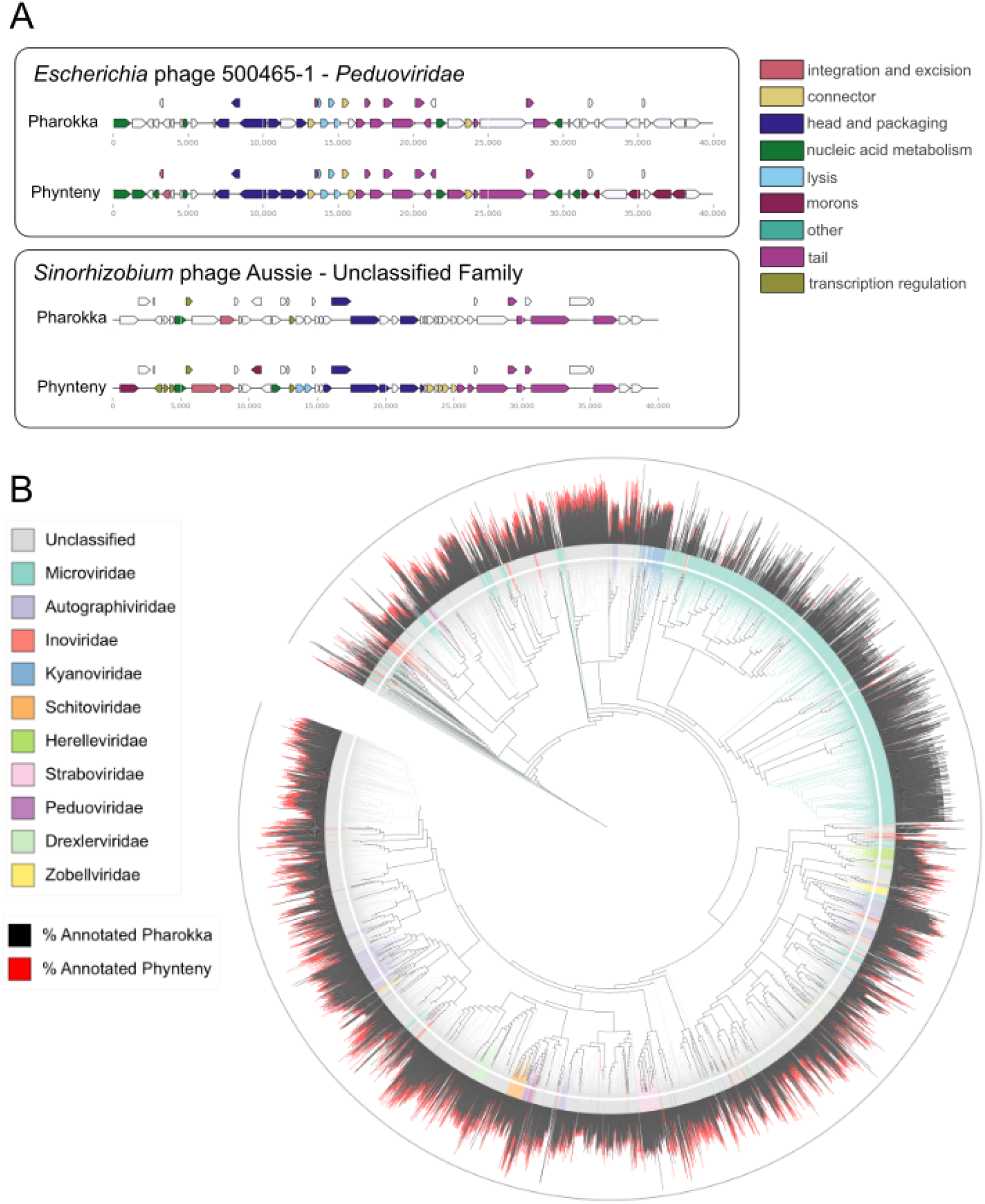
Phynteny increases the functional annotation rate across the phage taxonomic tree. **A**. Representative genomes annotated with Phynteny. Escherichia phage 500465-1 (GenBank accession number CP025900.1) and Sinorhizobium phage Aussie (OR786373.1). **B**. Phylogenetic tree of all phage genomes in the INPHARED database. Branches and ring colour correspond to the family. The bar chart displays the annotation rate before and after annotation with Phynteny. The line outside corresponds to an annotation rate of 100%.

Across INPHARED, the average proportion of genes annotated per genome increased from 41% with Pharokka alone to 48% following Phynteny annotation. The degree of improvement varied substantially across viral families. For example, Microvirus genomes (n = 8,606), which encode a median of six genes, showed a very slight average increase of 0.3% (Figure 4B). In contrast, substantial improvements were seen in larger genomes with more syntenic structure, such as those from *Steigviridae* (+17.6%, n = 399, 76-103 kb), *Duneviridae* (+15.9%, n = 46, 39-47 kb), and *Peduoviridae* (+14.2%, n = 52, 11-51-kb) (Fig S6). Phynteny improves the portion of genes annotated across diverse taxa, including Unclassified phages, which make up the majority of the INPHARED database, suggesting that Phynteny may enable more comprehensive annotation of undercharacterised lineages.

### Gene-order enhanced annotation is concordant with protein structure prediction

To further evaluate the reliability of Phynteny’s predictions, we compared its functional annotations to those derived from protein structural similarity for phage genomes in INPHARED (Fig. 5A). Structural annotations were obtained using Phold, which uses the ProstT5 protein language model^25^ to search for protein structural homologs using Foldseek^41^. Phynteny showed strong concordance with structure-based predictions across most PHROG functional categories. For genes involved in integration and excision, head and packaging, lysis, tail, and transcription regulation, the categories that were learnt best by Phynteny, >90% of Phynteny annotations aligned with the corresponding structural category. Moderate agreement was observed for genes assigned to the morons (71%) and other (80%) categories, which are more functionally diverse and less syntenically conserved.

**Figure 5.**
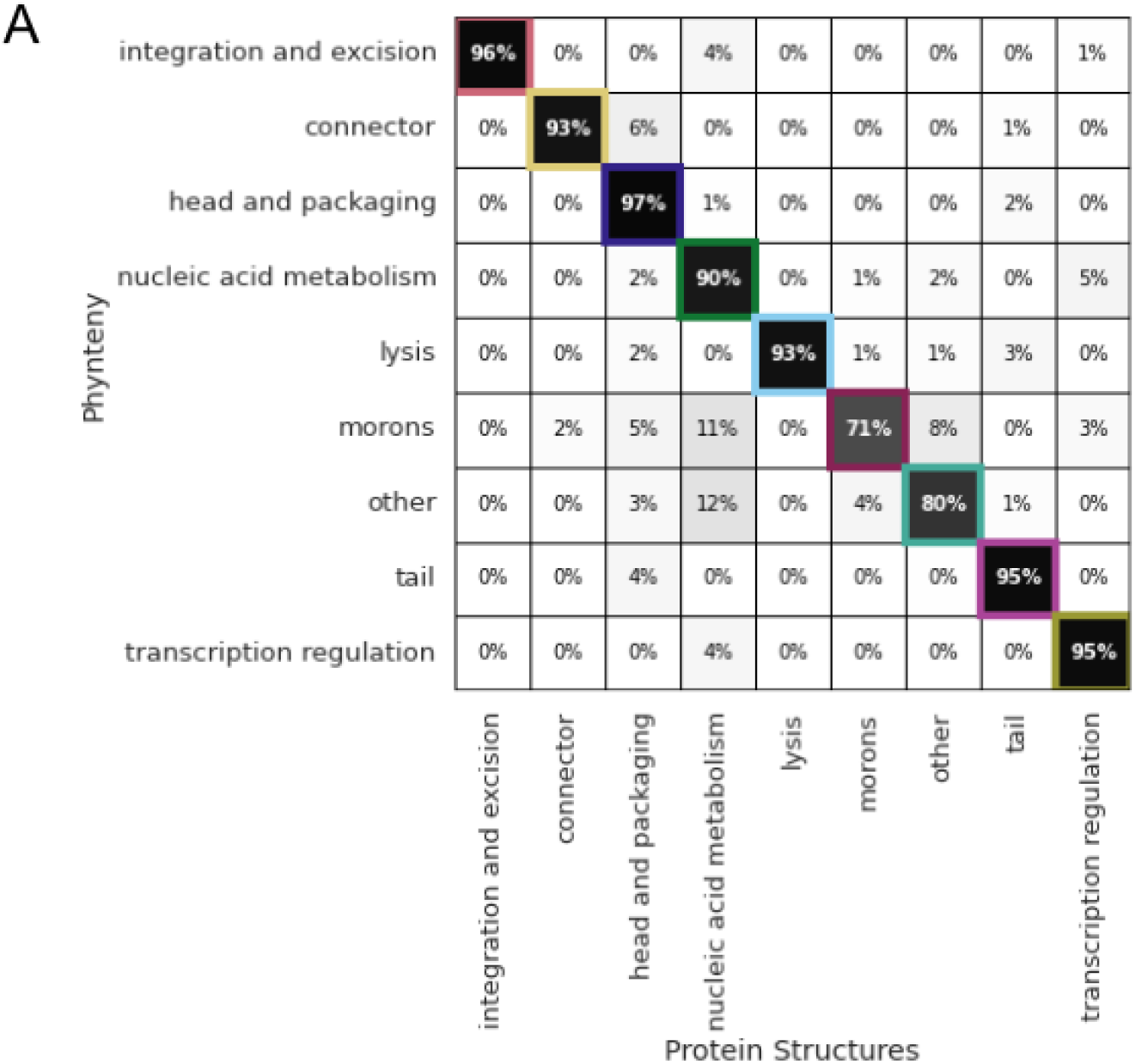
Agreement between Phynteny and structure-based function prediction. Confusion matrix of phage genomes in the INPHARED database annotated with Phynteny and structure-based function prediction.

To illustrate Phynteny’s ability to generalise functional assignments across diverse genomes, we examined the PHROGs most frequently annotated by Phynteny in the INPHARED database. Several of these PHROGs were consistently assigned functions by both Phynteny and structure-based prediction. For instance, phrog 600 (assigned to lysis), phrog 1989 (head and packaging), phrog 862 (nucleic acid metabolism), and phrog 13288 (tail) each appear in hundreds of genomes and consistently receive the same functional label across tools. Overall, Phynteny annotates 228 PHROG groups in more than 100 distinct genomes each (Tables S2 and S3), many of which previously lacked functional labels. These results demonstrate that Phynteny not only recovers accurate gene-level annotations but also provides functional resolution at the protein family level.

## Discussion

Gene order is a deeply conserved feature of phage genomes, maintained across diverse lineages due to physical constraints on DNA packaging and the need for coordinated gene expression during infection^26,27,42,43^. The gene ordering of phage genomes reflects functional groupings of modules for replication, structural proteins, and lysis, arranged in sequential blocks. Phynteny leverages this syntenic organisation to guide functional annotation of phage genes. While expert annotators have long used synteny as contextual evidence during manual annotation^30,44^, this reasoning is absent from automated phage genome annotation software.

Phynteny integrates gene-order information and amino acid composition using protein language model embeddings within a hybrid architecture that captures global gene order dependencies across phage genomes. It combines a bidirectional long short-term memory, which models local sequential relationships, with a Transformer that attends to broader contextual signals. To accommodate the circular topology of many phage genomes, Phynteny introduces circular attention, enabling the model to recognise relationships across genome boundaries. Protein embeddings are gradually introduced during training, encouraging the model to prioritise syntenic context before relying on sequence-level features. This architecture allows Phynteny to annotate a broader set of phage proteins, including those lacking detectable sequence homology, by leveraging the functional signals embedded in gene organisation, signals missed by traditional homology-based methods.

The annotation performance of Phynteny varies by functional category. Categories such as head and packaging, connector, and integration and excision exhibit the highest AUC values and prediction rates, likely due to strong conservation in both gene content and order. In contrast, categories like the monrons and other category are more sporadically distributed and less syntenically conserved, complicating prediction. This demonstrates that complementary methods may be used to achieve maximal functional annotation. The success of Phynteny also varies across taxonomic groups. Phages in the *Microviridae* family, which have compact genomes (∼5–7 kb) and between 3-14 genes and fewer conserved functional modules, show minimal gains^45,46^. Meanwhile, phages in the *Steigviridae*^*47,48*^, *Duneviridae*^*49*^ *and Peduoviridae*^*47,48*^ families, which have larger genomes, often share conserved genome architectures and well-characterised morphogenesis modules, show substantial improvements in annotation coverage.

To evaluate Phynteny’s predictive performance, we benchmarked its annotations against Phold, a structure-based annotation approach. Proteins that are unannotated using sequence homology but receive confident annotations from Phold serve as an ideal validation set as they are not homologous to the training data at the sequence level, nor are their annotations derived from sequence similarity. Phynteny effectively expands into these regions of the annotation space, complementing structure-based methods and demonstrating utility beyond classical homology inference. In practice, Phynteny can be applied after Phold to further increase annotation coverage. For example, many short genes (e.g., <200 amino acids) remain uncharacterised by structure-based annotation, since protein structure alone is often insufficient to predict the function of these small proteins^50,51^. Therefore, Phynteny can be applied after Phold to further increase annotation coverage, as implemented in the Sphae^20^ pipeline, where Phynteny is used to annotate genes that remain uncharacterised following structural analysis.

Phynteny assigns genes to one of nine broad functional categories, providing a general overview of genome organisation and functional potential. While coarse-grained, this classification enables consistent annotation across diverse phage genomes and facilitates interpretation of functional modules. However, many biologically relevant distinctions, such as between tail protein subtypes, depolymerases, antimicrobial resistance genes, and lysis mechanisms, are not yet captured. More detailed phage protein classification systems, such as those proposed in the Empathi framework^52^, demonstrate the feasibility and value of expanding functional labels for phage genes. Incorporating such granularity into Phynteny represents a promising direction for future development, which could enhance the resolution and biological relevance of automated phage genome annotation.

Beyond phages, a similar gene-order-based strategy could be applied to bacterial genomes. While global gene order is less conserved in bacteria than in phages, functionally related genes in bacteria often cluster together in operons or other conserved neighborhoods, providing valuable context for functional inference^53^. In fact, the notion of inferring gene function from conserved gene clusters in bacteria dates back decades^54^. Recent synteny-aware tools for bacteria that integrate protein language model embeddings with operon context significantly outperforms homology-based methods^31,55^, even identifying previously unrecognized toxin genes by exploiting conserved gene order^31^. This suggests that Phynteny’s synteny-guided framework could be scaled to improve functional predictions in bacterial genomes, capitalizing on the operon structure and co-evolved gene clusters characteristic of bacteria.

Importantly, Phynteny builds upon and benefits from existing annotations, with the capacity to evolve as databases expand and new gene functions are experimentally characterised. The model can be retrained to integrate these updates, continually refining its predictions. All steps in the Phynteny pipeline are open-source and accessible, supporting ongoing updates and community-driven improvement. Phynteny is already deployed in the Sphae phage-therapy workflow^20^, where it enables high-resolution functional annotation of phage genomes. This is particularly valuable in phage therapy applications, where understanding gene function is essential for evaluating safety by predicting phage integration and identifying accessory genes such as toxins and resistance factors. More broadly, Phynteny demonstrates that conserved gene order is a powerful and underutilised signal for functional annotation in phage genomes. By combining syntenic context with protein embeddings in an architecture tailored to phage genome structure, it expands the scope of genes that can be confidently annotated beyond sequence homology. In doing so, Phynteny broadens our understanding of phage genome content, enables the discovery of novel genes, supports biotechnological applications, and deepens insights into viral roles within microbial ecosystems.

## Acknowledgements

This work was supported with the assistance of resources and services from Flinders University using the DeepThought High Performance Cluster (https://doi.org/10.25957/FLINDERS.HPC.DEEPTHOUGHT) and Pawsey Supercomputing Research Centre, which is supported by the Australian Government. We also acknowledge the use of ChatGPT/OpenAI for proofreading the manuscript and GitHub Copilot for debugging code. SRG was supported by scholarships from the Australian Government Research Training Program and the Playford Trust. RAE was supported by awards from the Australian Research Council DP250103825 and FL250100019.

## Methods

### Data

#### Training Data

All phage genomes were retrieved from the PhageScope database (873,718 genomes)^9^. To mitigate redundancy, a single representative was selected from each PhageScope subcluster, defined bya threshold of identity >0.6 and coverage >0.75 (531,031 genomes). The quality of the genomes was determined using CheckV (version 1.0.3)^56^ and genomes with ‘low’ or ‘undetermined’ quality were removed (295,461 genomes remaining). Genomes were annotated with Pharokka (v.1.7.3)^14^ using pyrodigal-gv^57^ to predict coding sequences and mmseqs2^58^ to match the coding sequences to the PHROGs database^22^. Genomes that contained fewer than five genes predicted by pyrodigal-gv and with no genes annotated with a known PHROG category were removed. The remaining 289,106 genomes were used for training.

#### Prophage Validation Set

Prophage sequences extracted from all complete bacterial genomes in the NCBI assembly repository (downloaded 1st of June 2022) using Phispy (version 4.1.22)^59^. Prophage regions that were at least 5,000 bp in length, contained five or more genes, and contained at least one gene annotated as a phage gene were considered, yielding 5,005,011 high-quality prophages detected from 945,551 bacterial genomes^39^. Prophage sequences were clustered with the same procedure used by PhageScope^9^. Sequences were clustered using mmseqs easy-cluster (version 12.113e3)^58^ with a threshold of identity >0.9 and coverage >0.9 (1,523,202 clusters). Subsequently, the representative sequences of these clusters were used as the inputs to another round of clustering with a threshold of identity >0.6 and coverage >0.75, resulting in 947,791 clusters.

To eliminate overlap between the prophage sequences and the Phynteny training data, the prophage sequences were aligned to the training data using mmseqs easy-search (version 12.113e3) with a threshold identity of >0.5 and coverage of >0.3. The 696,058 prophage sequences, which aligned to the PhageScope training data, were removed from the validation data, resulting in 251,733 prophage sequences. The quality of the remaining sequences was evaluated using CheckV, and sequences with ‘low’ or ‘undetermined’ quality or a predicted contamination above 15% were removed, leaving 16,679 sequences.

Prophage sequences were annotated with Pharokka using pyrodigal-gv to predict coding sequences and mmseqs2 to match coding sequences to the PHROGs database. Sequences without at least one known PHROG category were removed, leaving 16,442 prophage sequences.

#### INPHARED Dataset

Taxonomic comparisons were made using the INfrastructure for a PHAge REference Database^11^ (April 2025), which includes all phage genomes deposited in NCBI GenBank along with associated taxonomy and host information where available. Compared to other resources such as PhageScope, INPHARED provides more comprehensive and standardised metadata.

Each genome was annotated using Pharokka^14^ using pyrodigal-gv to predict coding sequences and mmseqs2 to match coding sequence to the PHROGs database. Unknown genes were then annotated with Phynteny. To generate the INPHARED phylogenetic tree, genomes in the INPHARED database were de-replicated at 90% ANI with dRep^60^ (version 3.4.2) using the options: “—ignoreGenomeQuality -l 5000 -pa 0.90—SkipSecondary” resulting in 6,046 genomes. Genomes were filtered to remove genomes annotated by INPHARED as belonging to the ‘Unclassified’ family.

#### Encoding

Each phage genome was encoded as an x 1292 matrix, where denotes the number of protein-coding genes in the genome. Genes were ordered sequentially based on their genomic start positions. The first nine columns of the matrix captured functional annotations using one-hot encoding, with each column representing one of nine distinct PHROG qfunctional categories. Gene orientation was encoded in the subsequent two columns using a binary scheme to indicate positive (0) or negative (1) strand orientation. The following column encoded gene length as the number of nucleotides in the coding sequence, normalised by a factor of 1,000. The remaining 1,280 columns contained contextualised protein sequence embeddings generated using the ESM-2 language model^34^ generated using the esm2_t33_650M_UR50D.pt model using the HuggingFace transformers library (version 4.49.0)^61^.

#### Model Architecture and Training

The *Phynteny* model is constructed using a transformer-based architecture for the categorical classification of gene functions in phage genomes. The model integrates multiple components to capture both local and global dependencies within phage genomic contexts, including dedicated embedding layers, a BiLSTM module, a transformer encoder block, and a final classification head. To improve the model’s capacity to learn positional and contextual patterns, we additionally incorporated sinusoidal positional encodings and a circular relative attention mechanism. All components were implemented in PyTorch (version 2.6.0)^62^.

#### Dynamic masking

A dynamic masking process was implemented to address class imbalance during training and ensure a balanced representation of each PHROG category. Inverse frequency weights *w*__*i*__ are computed for each PHROG category *i* in the training dataset, where *f*__*i*__ is the frequency of a PHROG category in the dataset and normalised to sum to one such that:

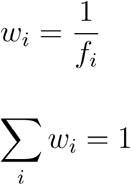

For each input sequence *x*, we define a probability distribution *q* over the genes with known PHROG category. This distribution is adjusted using the computed class weights to create a weighted probability distribution *q*__w__, ensuring that less frequent classes have a higher chance of being selected for masking:

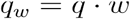

The number of tokens to mask is determined as a portion (default = 0.15) of the total maskable tokens, ensuring at least one token is masked. Tokens are then sampled based on the weighted probability distribution using a Bernoulli distribution, and the selected tokens are masked as unknown in PHROG category one-hot encoding. This dynamic masking process is performed at each epoch, introducing label variability across epochs. This promotes model generalisation and mitigates overfitting by preventing the memorisation of static annotation patterns.

#### Custom Masked Protein Dropout Layer

To ensure model generalisability, we developed a custom Masked Protein Dropout layer that selectively applies dropout to protein embeddings while preserving gene order information. This mechanism allows the transformer architecture to learn both gene order patterns and protein sequence features simultaneously.

Dropout is applied only to the protein embedding of masked tokens with probability *p*, where *p* represents the dropout probability, while gene order, functional category, gene orientation and length remain intact:

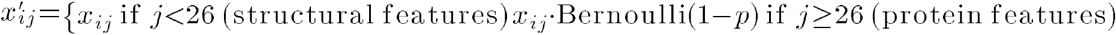

Where *x*__*ij*__ is the input embedding at position and feature dimension *j*. The value 26 represents *d*__*gene*__, the total dimensionality of gene structural features, calculated as the sum of:

- Function embedding dimension (16): Learned categorical embeddings representing gene functional classes
- Strand embedding dimension (2): Linear transformation of one-hot encoded gene orientation (forward/reverse strand)
- Length embedding dimension (8): Linear transformation of normalized gene length features

Additionally, we implemented a progressive dropout schedule that gradually reduces the dropout rate from an initial value *p*_*initial*_ (typically 1.0, complete masking) to a final value *p*_*final*_ of 0.7 over *n* epochs:

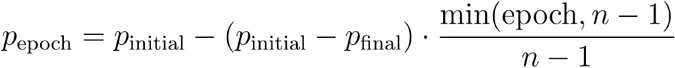

This progressive approach encourages the model to first learn gene order dependencies before incorporating protein sequence information, creating a curriculum learning effect that improves generalisation. During inference, the dropout layer is disabled, allowing the model to utilise all available features.

#### Embedding layers

To convert sparse, categorical representations into dense, continuous representations, the encoded phage genomes are input to the model using separate learnable embedding layers for the functional one-hot encoding (16 dimensions), strand embedding (2 dimensions), gene length (8 dimensions) and protein language model embedding (230 dimensions). The outputs of these embedding layers were concatenated to form the final input embedding for each sequence and then passed through the BiLSTM and transformer.

#### Sinusoidal Positional Encodings

To incorporate positional information into the model, we use sinusoidal positional embeddings ^38^. These embeddings are generated using sine and cosine functions of different frequencies, allowing the model to learn relative positions in sequences. The sinusoidal positional embedding for a given position *p^os^* and dimension *i* is defined as:

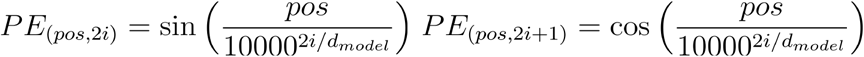

where *d*_*model*_ is the dimension of the model.

These embeddings have several desirable properties: they can handle variable-length sequences, provide a unique encoding for each position, and allow the model to extrapolate to sequence lengths not seen during training. The sinusoidal functions with different frequencies create a distinct pattern for each position while maintaining a smooth relationship between nearby positions, which helps the model learn relative positional relationships.

#### Bidirectional LSTM

A BiLSTM layer captures sequential dependencies in both forward and backward directions. The BiLSTM layer processes the concatenated embeddings and outputs a hidden representation that is fed into the transformer encoder.

#### Circular relative attention

As most phage genomes are circular, we introduce a circular relative attention mechanism to capture the relative positions of tokens in a circular manner. This mechanism computes attention scores by considering both direct and wrap-around distances between tokens.

In the transformer attention mechanism, each input token is transformed into three representations^38^:

- Query (Q): represents the information that each token seeks from other tokens in the sequence.
- Key (K): represents the information that each token advertises or makes available to other tokens. V
- Value (V): represents the actual information content that is retrieved and passed forward when a token receives attention.

These are computed as linear transformations of the input embeddings:

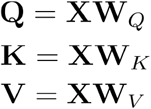

where X is the input sequence and *W*_*Q*_, *W*_*k*_,*W*_*v*_ are learned parameter matrices.

Our circular relative attention mechanism extends the standard attention computation by incorporating learnable relative position encodings that account for the circular nature of phage genomes. The relative position encodings for keys and values are learnable parameters initialised randomly, enabling the model to learn optimal representations for circular positional relationships.

The attention mechanism is defined as:

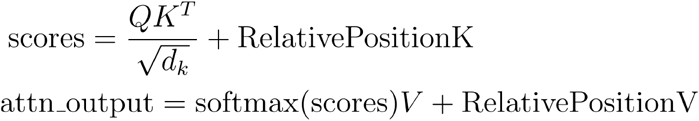

where Q, K, and V are the query, key, and value matrices respectively, d_k is the dimension of the keys (used for scaling to prevent vanishing gradients), RelativePositionK captures circular positional relationships between queries and keys, and RelativePositionV modifies the value representations based on circular relative positions. This formulation allows the model to effectively handle circular sequences by learning position-aware attention patterns that can span across the genome boundaries.

Transformer Encoder and Classification Head

The transformer encoder consists of two custom transformer encoder layers, which incorporate the circular relative attention mechanism. Each layer is followed by a feedforward neural network and layer normalisation. The final classification head is a fully connected linear layer that maps the output of the transformer encoder to the desired number of output classes. Standard dropout is applied to the output of the transformer encoder to prevent overfitting.

#### Model Optimization

The model is optimized using the AdamW optimizerr^63^ with a weight decay of 0.01, which helps prevent overfitting by penalizing large weights. The learning rate follows a cosine annealing schedule with linear warmup:

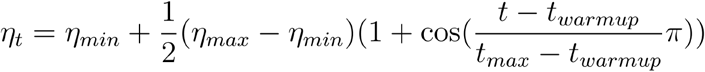

for, *t> t_warmup_*, and a linear warmup phase:

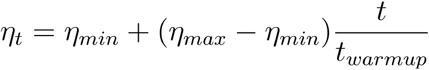

for *t* ≤*t _warmup_* where *t* is the current training step, *t _warmup_* is the number of warmup steps, and is the total number of training steps.

The initial learning rate (η*_max_*) is set to 5e-4, with a minimum learning rate (η*_min_*) of 5e-6. The warmup period spans the first 5% of training steps, allowing the model to establish stable gradient updates before applying the full learning rate. Gradient clipping is applied with a maximum norm of 1.0 to prevent exploding gradients.

#### Diagonal Loss Function

The primary loss function is categorical cross-entropy, defined as:

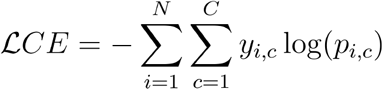

where *N* is the batch size, *C* is the number of functional classes, *y*_*i,c*_ is the ground truth label, and *p*_*i,c*_ is the predicted probability.

Additionally, a custom diagonal loss function is used to encourage the model to focus on non-diagonal elements in the attention matrix, which aids in capturing long-range dependencies:

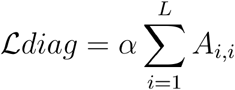

where *A* is the attention weight matrix, *L* is the sequence length, and α is a scaling factor. The total loss is then computed as:

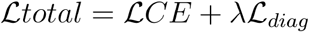

Where λ controls the contribution of the diagonal loss. This combined loss function penalizes the model when attention weights are concentrated along the diagonal, promoting a more distributed attention pattern that better captures relationships between different genes in the phage genome.

#### Training

Phynteny was trained using ten-fold cross-validation using the scikit-learn (version 1.5.1) *Kfold* method^64^ with the parameters mask_portion=0.15, learning_rate=10^-4^, hidden_dimension=512, num_transformer_layers=2, batch_size=16, protein_dropout_rate 0.7 for a total of 50 epochs (Fig. S7). During training, the AdamW optimiser is used to update the model parameters, benefiting from weight decay regularisation to prevent overfitting. A cosine learning rate scheduler with warmup is employed to optimise the learning rate dynamically.

Training runs were completed on a single AMD EPYC 7A53 “Trento” 64-Core GPU Node with eight AMD Instinct MI250X GPUs and 256 GB RAM provided by the Pawsey Supercomputing Resource Centre. We performed a grid search over key hyperparameters (e.g., learning rate, hidden dimension, dropout). Multiple configurations yielded comparable performance, suggesting robustness to hyperparameter variation (Table S5).

#### Confidence Calibration

Predictions are generated using an ensemble of all ten cross-validation models. To combine their outputs, we compute a Phynteny score for each gene prediction. This score is obtained by summing the softmax vectors from the ten models, yielding a score for each PHROG category, with a maximum possible value of ten. The predicted gene function is assigned to the category with the highest Phynteny score. The score is not normalised to a 0–1 scale to avoid misinterpretation as a calibrated confidence measure. This approach is analogous to the PhANNs score used in PhANNs^23^.

To provide well-calibrated confidence scores for each functional prediction, we employed isotonic regression as a post-processing step on the model’s raw output probabilities using the *sci-kit* learn isotonic regression method. For each functional category, we collected the model’s Phynteny scores and the corresponding binary ground-truth labels (correct/incorrect). We then fit an isotonic regression model *f*_*c*_ for each category *c* using the training data. The isotonic regression model learns a monotonically increasing function that best maps the raw scores to the observed frequencies of correct predictions.

For a new prediction with Phynteny score *s* in category *c*, the calibrated confidence is given by:

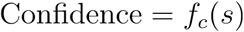

where *f*_*c*_ is the isotonic regression model for category *c*. This approach ensures that the confidence scores are interpretable as true probabilities and are distributed across the full [0,1] range. Calibration quality was assessed using the Brier score and log loss, confirming that isotonic regression improved the reliability of the confidence estimates.

#### Validation, Performance Evaluation and Benchmarking

The performance of each model within the ensemble was evaluated on its respective validation dataset. Receiver operating characteristic (ROC) curves with the area under the curve (AUC) and precision-recall (PR) curves with the average precision score (APS) were calculated for each model in the ensemble using *scikit-learn*^*64*^ (version 1.2.2). These ROC and PR curves were generated on structural annotations generated using Phold (https://github.com/gbouras13/phold). As precision-recall curves produce lower than expected area under the curve for imbalanced datasets, Phold annotations were undersampled to ensure an equal number of gene predictions for each category.

Confusion matrices were generated using the *scikit-learn* confusion_matrix method.

#### Visualisation

The summed attention weights were visualised by summing the weights across attention heads. Attention between categories was visualised as a graph using Networkx (version 3.3) (Hagberg, Schult, and Swart 2008) for modelling the first training fold and its associated validation data.

ESM2 embeddings were visualised using pairwise controlled manifold approximation (PACMAP) dimension reduction using the pacmap package (version 0.8.0)^65^.

To visualise the improvement in annotation rate provided by Phynteny across phage families, phage genomes in INPHARED were dereplicated at 90% average nucleotide identity with dRep (version 3.6.2; options: dereplicate --ignoreGenomeQuality -p 32 -l 5000 -pa 394 0.90–SkipSecondary)^60^ which resulted in 6,046 representative genomes. Representative genomes were used to generate a phenetic tree using mashtree (version 1.4.3)^66^ using default parameters and visualised using iTOL^67^ and coloured according to family designation via INPHARED.

Figures of phage genomes were generated using pyGenomeViz (version 1.5.0)(https://github.com/moshi4/pyGenomeViz).

## Data availability

Pretrained Phynteny models are available at https://doi.org/10.5281/zenodo.15276213. Prophage sequences identified using Phispy from all prophage genomes are available in fasta format at https://doi.org/10.25451/flinders.22317060.v1 and in GenBank format at https://doi.org/10.25451/flinders.22317059.v1. Additional details of these genomes can be downloaded from https://doi.org/10.25451/flinders.22299673.v1.

## Author Contributions

SRG, PD and RE conceived the project. SRG wrote the paper, designed models and datasets, performed the analysis and performance evaluation and developed the software. GB, BP, VM and MR providence input for the analysis and the software. BP was responsible for implementing Phynteny in the Sphae pipeline. RAE supervised the project. All authors reviewed and approved the paper.

## Code availability

Phynteny is an open-source software, and its code can be found at https://github.com/susiegriggo/Phynteny_transformer/ and is available on bioconda https://anaconda.org/bioconda/phynteny_transformer and PyPi https://pypi.org/project/phynteny-transformer/. Python scripts used to train Phynteny models is available at https://github.com/susiegriggo/Phynteny_transformer/tree/main/train_transformer. Analysis code used to generate figures is available at https://github.com/susiegriggo/Phynteny_transformer/tree/main/notebooks.

