## Supplementary Information for "Synteny-aware functional annotation of bacteriophage genomes with Phynteny"

### Supplementary Figures

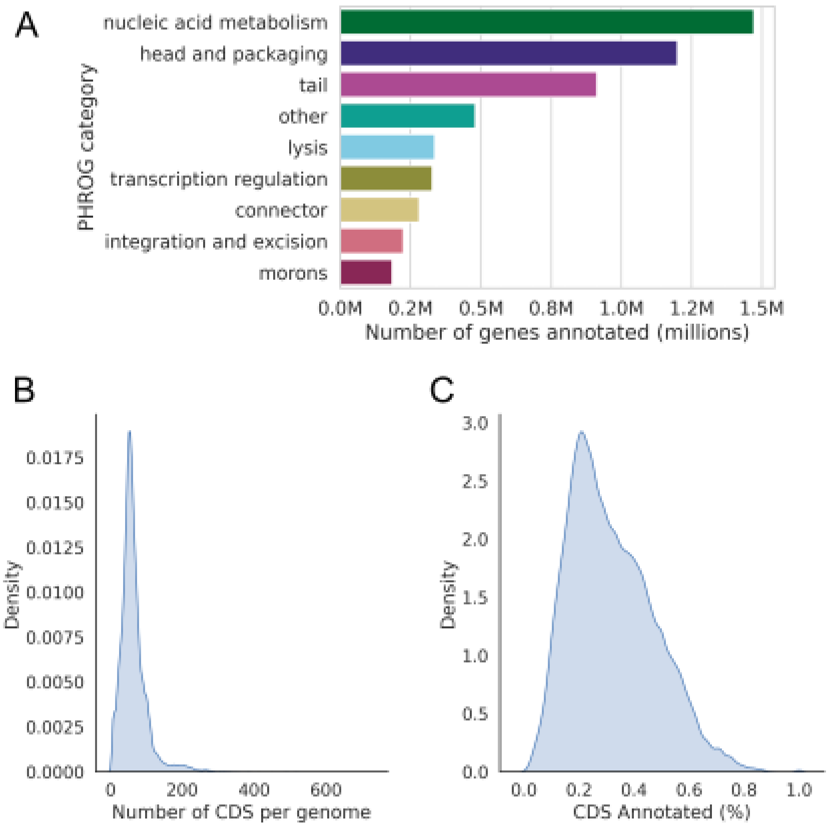


**Fig S1: PHROG annotation coverage of bacteriophage genes in the PhageScope dataset.**

A. Number of annotated genes assigned to each of the nine PHROG functional categories across all genomes. The total number of genes identified was 18,414,322.
B. Distribution of the number of genes per genome (289,106 genomes).
C. Distribution of the percentage of genes annotated with PHROGs per genome (289,106 genomes).

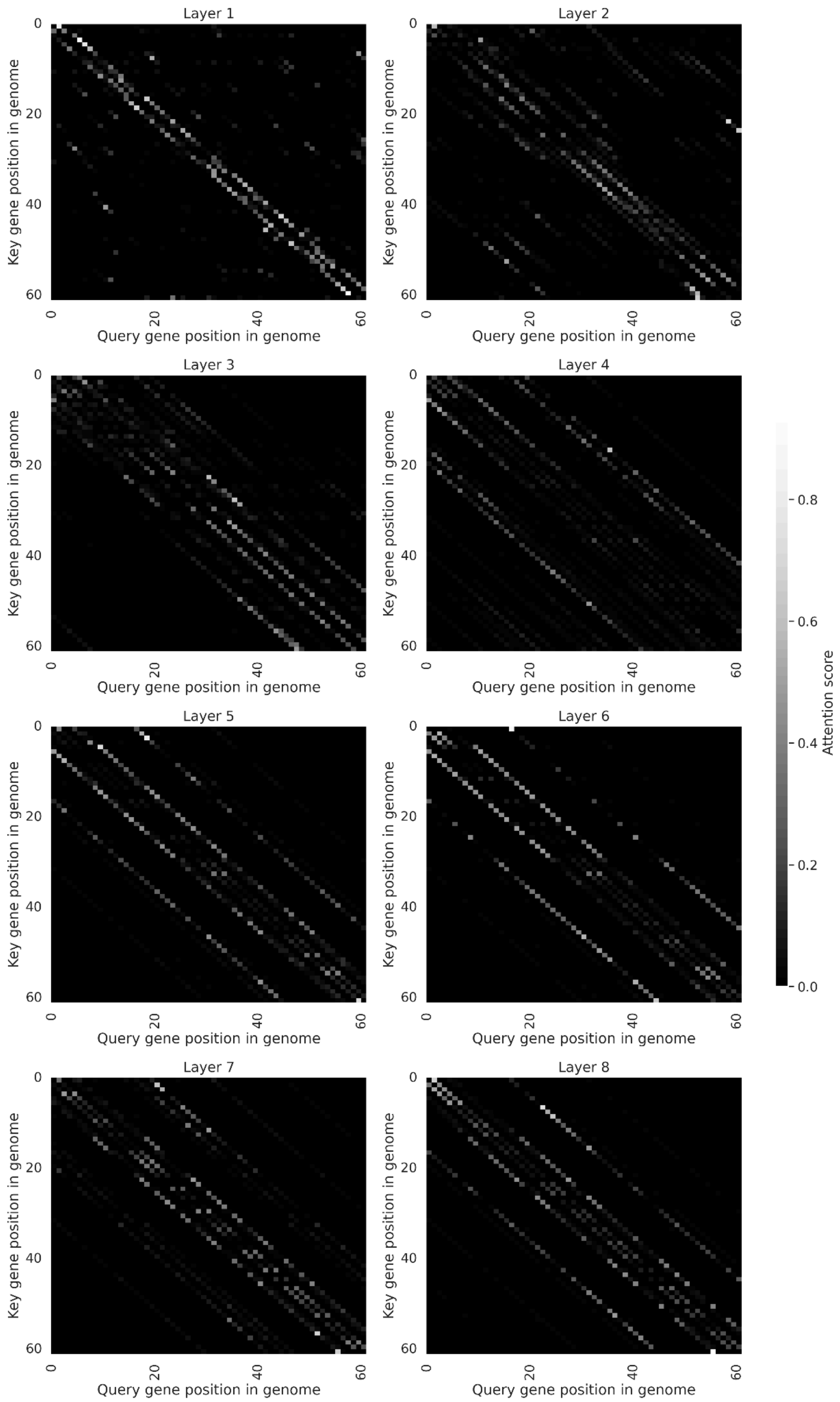


**Fig S2: Attention weights across eight transformer layers for a single phage genome.** Self-attention maps from a representative genome (Geobacillus phage phiOH2; AB823818), showing the attention weights across each transformer layer (8 layers) of the first model in the Phynteny ensemble. The attention weights range from 0 (black) to 1 (white). The white lines near the diagonal indicate that the model is attending to neighbouring genes, while the more dispersed, lighter points indicate that the model is attending to distant genomic regions. The model attends both to neighbouring genes and distant genomic regions, reflecting context-aware information aggregation.

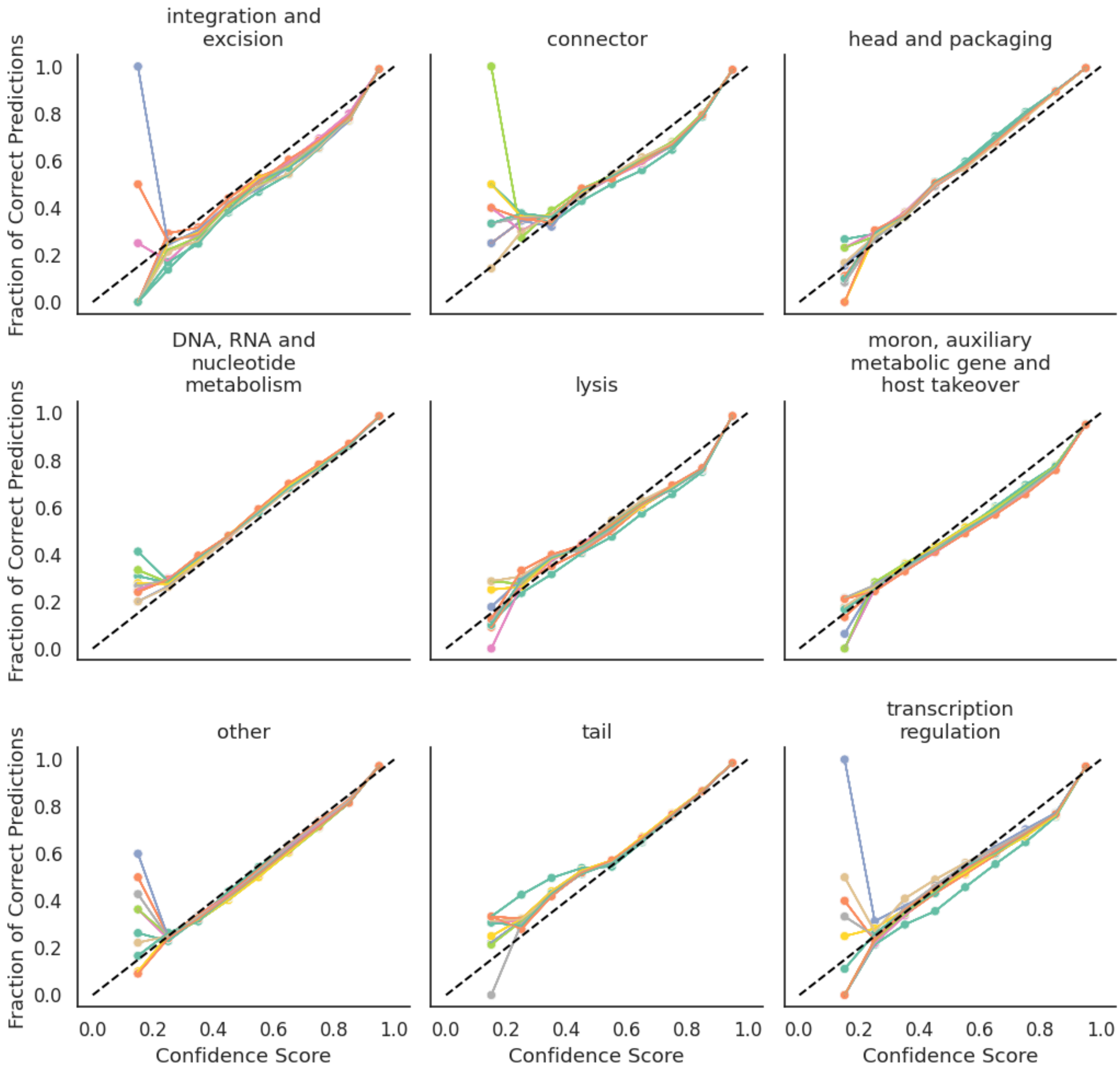


**Fig S3: Confidence calibration for each category in Phynteny Transformer.** Each line represents a model obtained from 10-fold cross-validation.

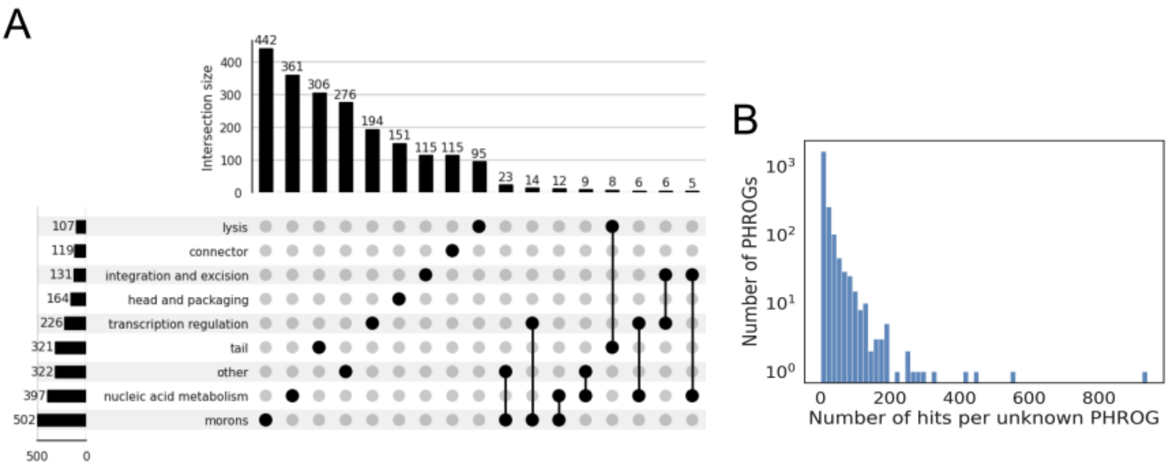


**Figure S4. Overlap of functional categories and distribution of annotation counts for unknown PHROGs annotated by Phynteny for the prophage dataset.**

A. UpSet plot showing intersections between PHROG functional categories predicted by Phynteny. PHROGs that are annotated by Phynteny only once are ignored. Only intersections with at least five PHROGs are shown.

B. Distribution of the number of hits per PHROG, excluding PHROGs annotated by Phynteny only once. The y-axis is shown on a logarithmic scale.

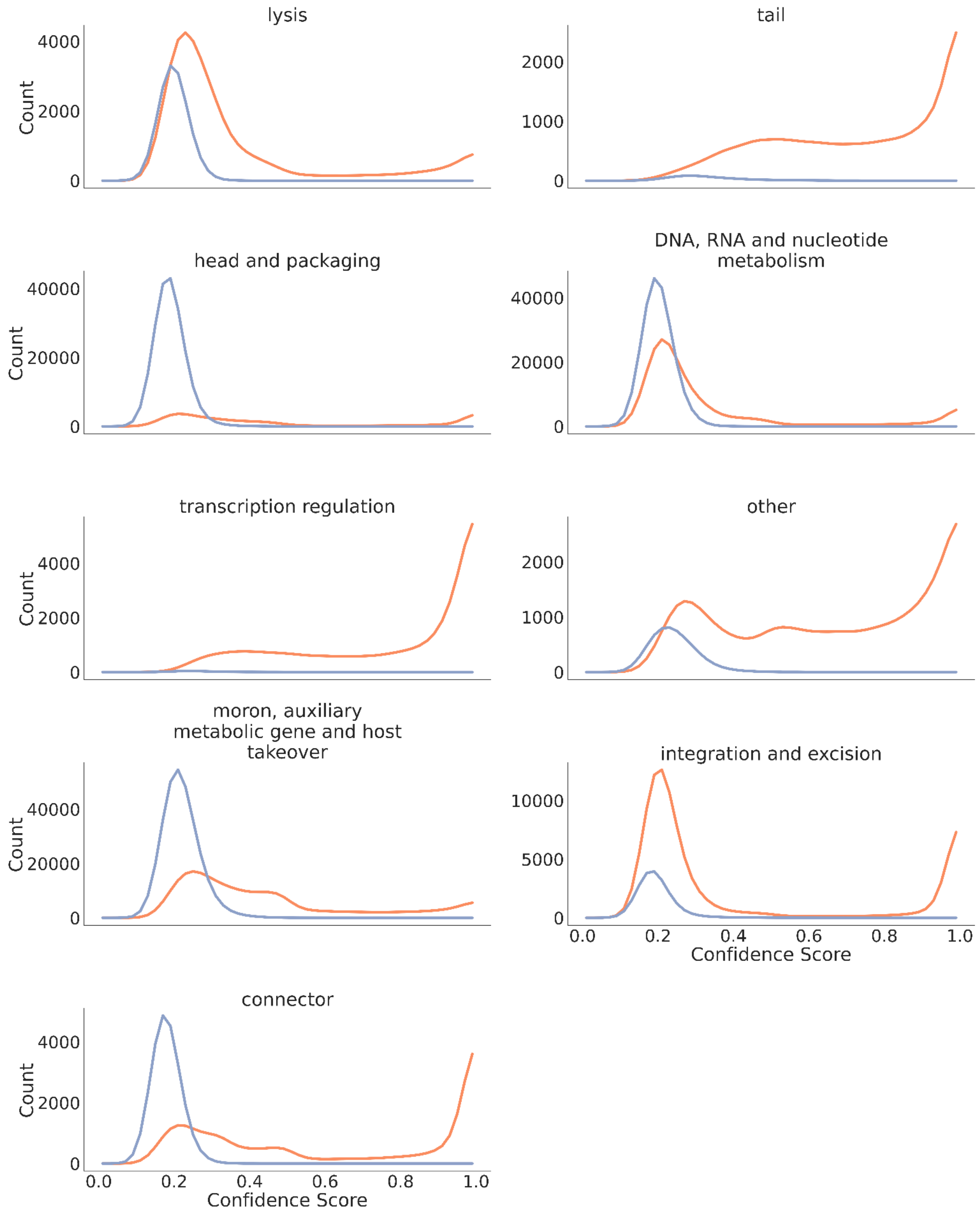


**Fig S5: Distribution of confidence scores for Phynteny annotations on unknown prophage proteins across nine functional categories using different Phynteny architectures.** The orange curves represent predictions from the complete Phynteny model, which incorporates Transformer and LSTM layers, while the blue curves show results from a simplified model trained without these sequence-contextual layers. For each functional category, the removal of Transformer and LSTM components resulted in a shift towards lower confidence scores.

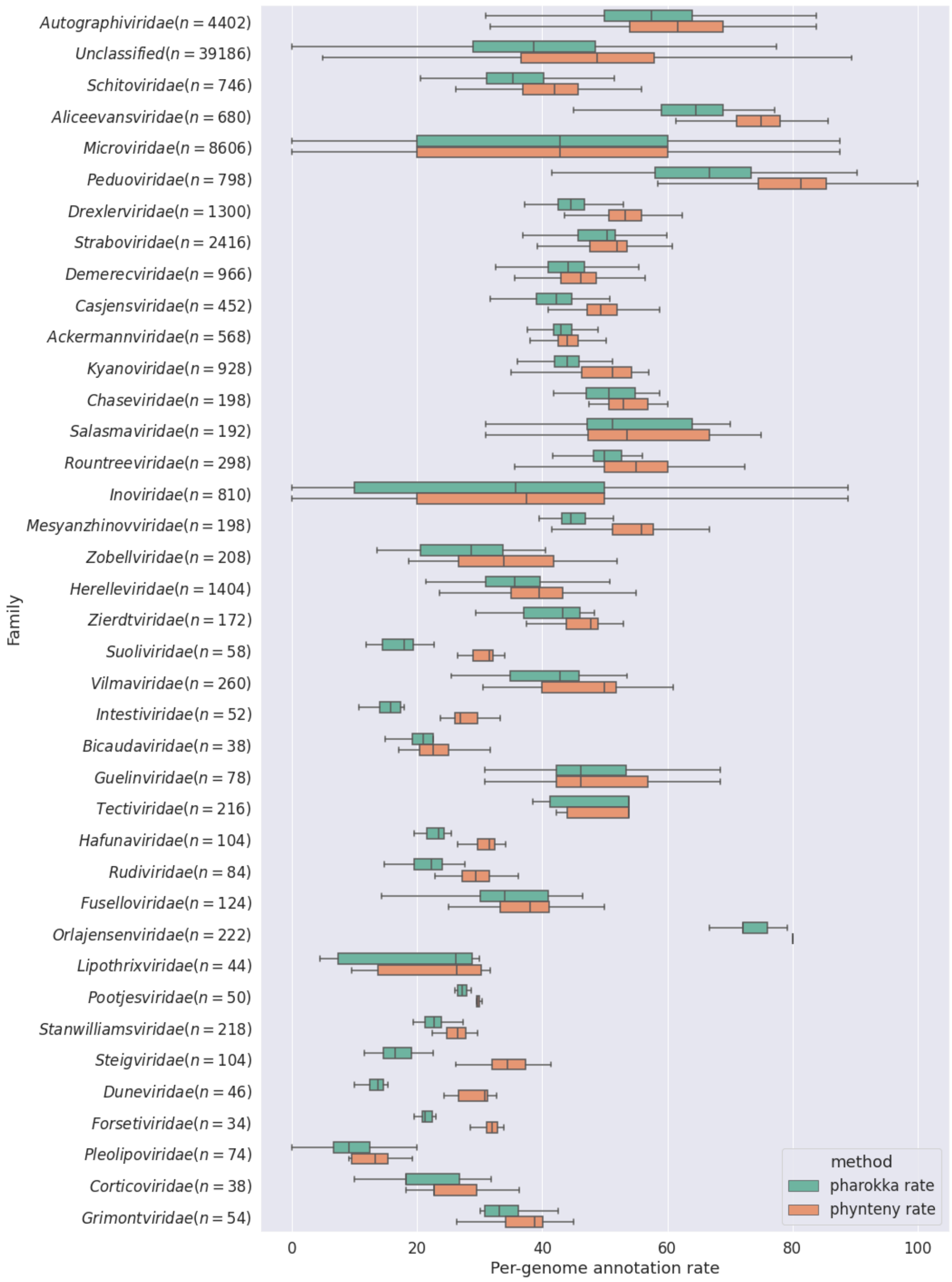


**Fig S6: Comparison of per-genome annotation rates across families.**
Distribution of annotation rates for annotation with pharokka only and pharokka followed by phynteny for INPHARED genomes. Only families with ≥15 genomes are shown.

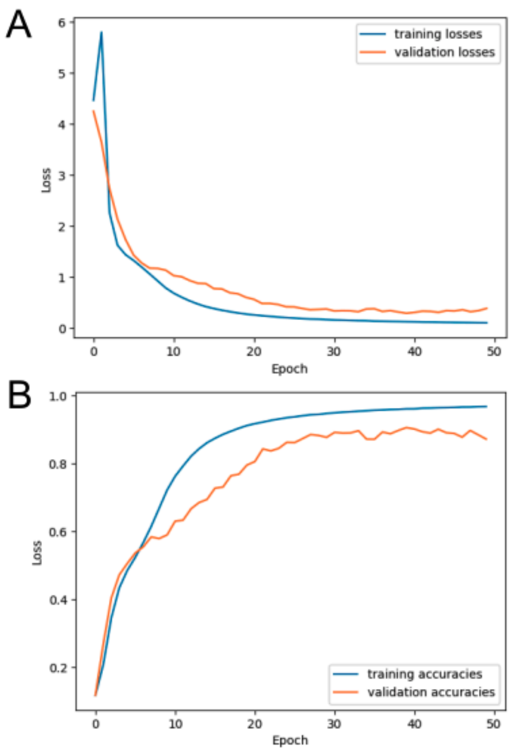


**Fig S7: Training and validation performance over 50 epochs of the Phynteny Transformer model.**

A. Training and validation loss.
B. Training and validation accuracy.

**Supplementary Tables**

**Table S1:** Brier score and log-loss of isotonic regression for Phynteny calibration.

| **category** | **Brier score raw** | **Brier score calibrated** | **Log loss raw** | **Log loss calibrated** |
|---|---|---|---|---|
| integration and excision | 0.003476 | 0.003296 | 0.017489 | 0.014617 |
| connector | 0.004052 | 0.003712 | 0.017293 | 0.015345 |
| head and packaging | 0.015812 | 0.015384 | 0.06363 | 0.058233 |
| nucleic acid metabolism | 0.016718 | 0.016363 | 0.07086 | 0.063977 |
| lysis | 0.005959 | 0.005657 | 0.028985 | 0.024763 |
| morons | 0.011823 | 0.01131 | 0.054333 | 0.045477 |
| other | 0.013275 | 0.0129 | 0.056904 | 0.051065 |
| tail | 0.013139 | 0.012817 | 0.051627 | 0.048274 |
| transcription regulation | 0.006077 | 0.005376 | 0.029537 | 0.023502 |

**Table S2:** Per-category Phynteny hits for phrog groups with more than 100 Phynteny predictions with confidence >0.8 in the INPHARED dataset.

*See the separate spreadsheet "Supplementary Tables".*

**Table S3**: Per-category Phold hits for phrog groups with more than 100 Phynteny predictions with confidence >0.8 in the INPHARED dataset.

*See the separate spreadsheet "Supplementary Tables".*

**Table S4:** Hyperparameters and training and validation loss for the transformer model trained with different configurations. The table summarises the tested configurations of the transformer model, including batch size, masking portion, learning rate (lr), dropout rate, hidden dimension size (hidden_dim), number of attention heads (num_heads), number of layers (num_layers), and protein-specific dropout (protein_dropout). For each configuration, the table reports the maximum training and validation accuracies, as well as the minimum training and validation losses achieved during the training process.

*See the separate spreadsheet "Supplementary Tables".*
